# Knocking out *SOBIR1* in *Nicotiana benthamiana* abolishes functionality of transgenic receptor-like protein Cf-4

**DOI:** 10.1101/2020.08.25.263483

**Authors:** Wen R.H. Huang, Christiaan Schol, Sergio Landeo Villanueva, Renze Heidstra, Matthieu H.A.J. Joosten

**Affiliations:** Laboratory of Phytopathology, Wageningen University, Droevendaalsesteeg 1, 6708 PB Wageningen, the Netherlands; Laboratory of Molecular Biology, Wageningen University, Droevendaalsesteeg 1, 6708 PB Wageningen, the Netherlands

## Abstract

The first layer of plant immunity is formed by pattern recognition receptors (PRRs) that are present at the cell surface and perceive extracellular immunogenic patterns. Receptor-like proteins (RLPs), such as the tomato (*Solanum lycopersicum*) PRR Cf-4 that provides resistance to the fungus *Cladosporium fulvum* secreting the matching avirulence factor Avr4, have an extracellular receptor domain consisting of leucine-rich repeats, but lack a cytoplasmic kinase domain for downstream signaling. RLPs constitutively interact with the receptor-like kinase SUPPRESSOR OF BIR1-1 (SOBIR1), thereby providing the receptor with a kinase domain, and recruit the co-receptor BRI-ASSOCIATED KINASE 1 (BAK1) upon their activation by a matching ligand. Trans-phosphorylation events, which can take place between the kinase domains of SOBIR1 and BAK1 after their association with the RLP, are thought to initiate downstream defense signaling. Currently, our knowledge on RLP/SOBIR1/BAK1-mediated defence initiation is limited and to understand the role of SOBIR1 in RLP function, we knocked out *SOBIR1* and its close homolog *SOBIR1-like* in the model plant *Nicotiana benthamiana*, as well as in transgenic *N. benthamiana* stably expressing *Cf-4*. We observed that Cf-4 function is completely abolished in the knock-out mutants, and we show that these plants can be used to perform transient complementation studies with SOBIR1 mutants. Thereby, these mutants are an important tool to study the fundamentals of plant immunity mediated by RLPs.

---

Plants are continuously challenged by a plethora of agents causing biotic stress. To defend themselves, plants have developed a multi-layered immune system (Jones and Dangl, 2006; Couto and Zipfel, 2016). The first layer is mediated by pattern recognition receptors (PRRs) that localize on the plasma membrane and perceive extracellular immunogenic patterns (ExIPs) (Dodds and Rathjen, 2010; Gust et al., 2017; van der Burgh and Joosten, 2019). So far, all known plant PRRs that carry an extracellular receptor domain consisting of leucine-rich repeats (LRRs) are either receptor-like kinases (RLKs) or receptor-like proteins (RLPs). They share the same overall structure, however, in contrast to RLKs, RLPs lack a cytoplasmic domain for downstream signaling (Liebrand et al., 2014; Couto and Zipfel, 2016). RLPs, such as the tomato (*Solanum lycopersicum, Sl*) PRR Cf-4 (Thomas et al., 1997) that mediates resistance against strains of the pathogenic extracellular fungus *Cladosporium fulvum* secreting the matching avirulence factor Avr4 (Joosten et al., 1994), have been shown to constitutively interact with the RLK SUPPRESSOR OF BIR1-1 (SOBIR1) (Gao et al., 2009; Liebrand et al., 2013). SOBIR1 is essential for Cf-4 accumulation and function (Liebrand et al., 2013), and upon recognition of Avr4 by Cf-4, the RLK BRI-ASSOCIATED KINASE 1 (BAK1), which is a regulatory co-receptor involved in development and defense, is recruited by the activated Cf-4/SOBIR1 complex (Postma et al., 2016). It has been proposed that subsequent trans-phosphorylation events between the kinase domains of SOBIR1 and BAK1 eventually initiate downstream defense signaling (van der Burgh et al., 2019). Although important advances have been made in deciphering PRR-mediated downstream immune signaling (Couto and Zipfel, 2016; van der Burgh and Joosten, 2019), little is known about how the RLP/SOBIR1/BAK1 complex exactly functions at the level of complex formation and downstream signal initiation. Here, we took advantage of the CRISPR/Cas9 system to knock out *SOBIR1* and its close homolog *SOBIR1-like* in the model plant *Nicotiana benthamiana* (*Nb*), as well as in *N. benthamiana* stably expressing the *Cf-4* transgene. Cf-4 is functional in *N. benthamiana* (Gabriëls et al., 2006), and here we demonstrate that Cf-4 function is completely abolished in an *N. benthamiana:Cf-4 sobir1*/*sobir1-like* knock-out mutant. We anticipate that these mutant lines are important materials for studying the fundamentals of plant immunity mediated by RLPs.

SOBIR1 is a positive regulator of plant immunity, as overexpression of *Arabidopsis thaliana* (*At*) *SOBIR1* in Arabidopsis, as well as in *N. benthamiana*, leads to constitutive activation of cell death and defense responses (Gao et al., 2009; Wu et al., 2017; van der Burgh et al., 2019). Surprisingly, no symptoms of constitutive immunity were observed when tomato *SlSOBIR1* or *NbSOBIR1* was overexpressed in *N. benthamiana* (Wu et al., 2017). Therefore, over the past years, the *At*SOBIR1-induced constitutive immunity in *N. benthamiana*, visible as a hypersensitive response (HR) at the site of agro-infiltration (transient expression) of *At*SOBIR1, is commonly employed to decipher the mechanism behind RLP/SOBIR1-mediated plant immunity. For this, endogenous *NbSOBIR1*(*-like*) are silenced in *N. benthamiana:Cf-4* by virus-induced gene silencing (VIGS), and subsequent complementation studies are performed by transiently expressing *At*SOBIR1 and various mutants of this RLK (Liebrand et al., 2013; van der Burgh, 2018). However, as VIGS only generates a gene knock-down, such complementation experiments require high amounts of repetitions due to variation caused by the presence of varying background levels that remain of the endogenous *Nb*SOBIR1(-like) protein. With the advent of the CRISPR/Cas9 gene-editing system, it is now possible to generate stable gene knock-outs in various plant species (Belhaj et al., 2013 and 2015; Ran et al., 2013), and this development prompted us to knock out functional *SOBIR1* in *N. benthamiana* and *N. benthamiana:Cf-4*, thereby generating a perfect system for complementation studies with mutants of SOBIR1.

To knock out both *SOBIR1* and *SOBIR1-like* in *N. benthamiana*, 6 single-guide RNAs (sgRNAs) targeting the open reading frames (ORFs) of both genes (Supplemental Figure S1 and Table S1) were designed using CRISPR-P 2.0 (Liu et al., 2017). Together with Cas9 and the selection marker BIALAPHOS RESISTANCE (BAR), sgRNA1, 2, 5, and 6 were assembled into the acceptor backbone (pAGM4723), referred to as Construct 1. Besides, sgRNA3, 4, 5, and 6 were cloned into the same acceptor backbone, referred to as Construct 2. The effectiveness and efficiency of the generated constructs were confirmed by studies based on their transient expression (Supplemental Figure S2), before stable transformation to explants of *N. benthamiana*. CRISPR/Cas9-induced mutations were detected by amplifying and sequencing the targeted gene regions, using isolated genomic DNA of the generated transformants as a template. Two homozygous *sobir1/sobir1-*like knock-out lines, generated by Construct 2, were obtained in the T1 generation (Supplemental Figure S3). *N. benthamiana sobir1/sobir1-like* line #1 contains a 1 bp deletion in the sgRNA3-target region and a 6 bp deletion in the sgRNA4-target region in the ORF of *SOBIR1*, whereas a 1 bp insertion is present in the ORF of *SOBIR1-like* (Figure 1A). Similarly, there is a 1 bp insertion and a 1 bp deletion in *SOBIR1*, and a 4 bp deletion in *SOBIR1-like* in the *N. benthamiana sobir1/sobir1-like* line #2 (Figure 1A).

**Figure 1.**
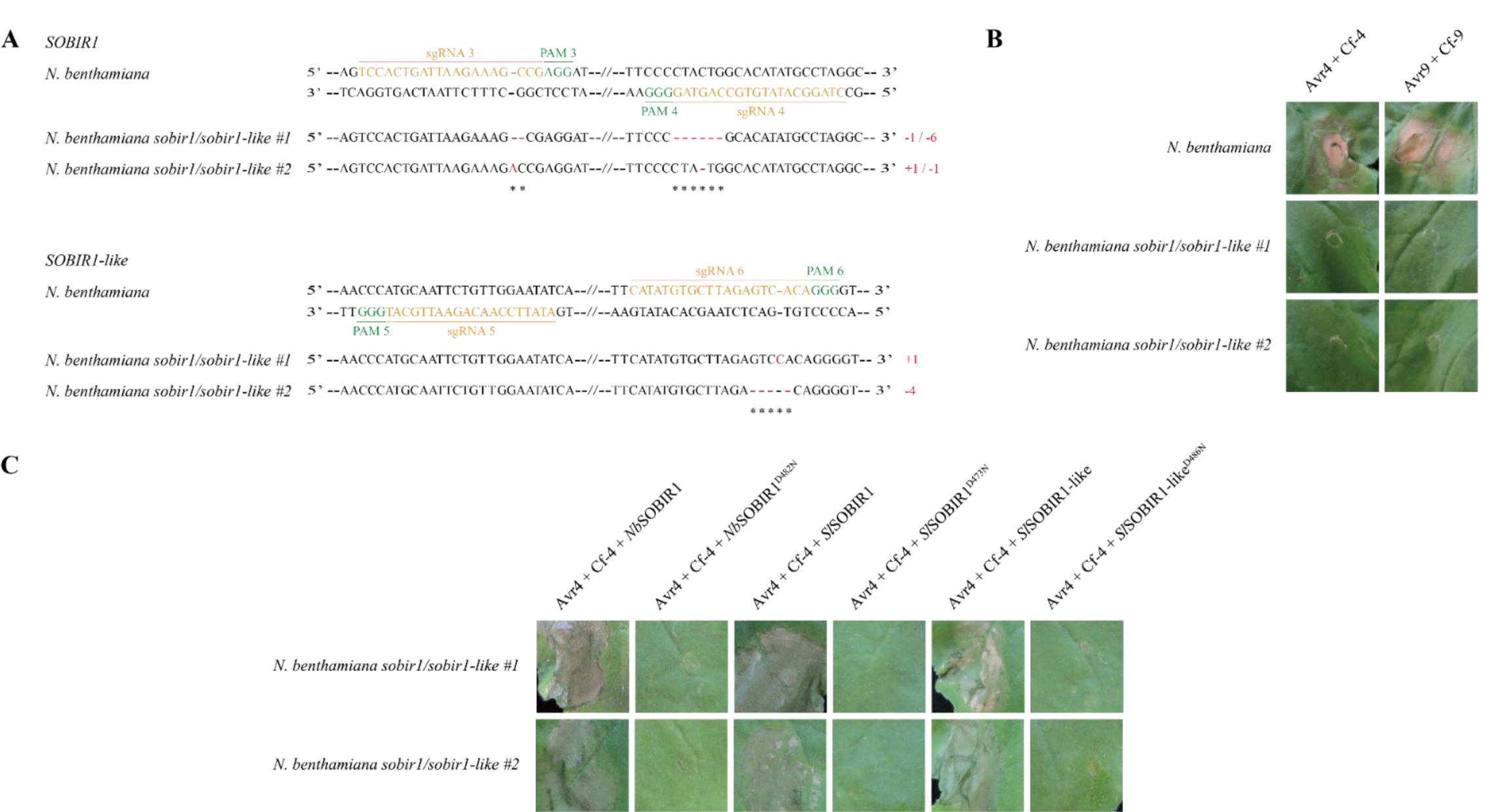
CRISPR/Cas9-induced targeted knockout of *SOBIR1* and *SOBIR1-like* in *N. benthamiana* abolishes the responsiveness to matching Avr/Cf combinations. A, Nucleotide sequence alignment of the regions in *SOBIR1* (upper panel) and *SOBIR1-like* (lower panel), targeted by single-guide RNAs (sgRNAs), with wild-type *SOBIR1* and *SOBIR1-like* sequences, respectively. The sgRNA sequences are indicated in orange, and the protospacer associated motifs (PAMs) are indicated in green. The deleted nucleotides in the generated transformants are indicated with red dashes, and the inserted nucleotides are denoted with red letters. The type of mutations and the numbers of deleted/inserted nucleotides are shown on the right. B, Transient co-expression of Cf-4 with the matching *C. fulvum* effector Avr4, or of Cf-9 with its matching *C. fulvum* effector Avr9, by *Agrobacterium*-mediated transient expression, triggers a rapid HR in the leaves of wild-type *N. benthamiana* plants (upper two panels), whereas neither Avr4/Cf-4-, nor Avr9/Cf-9-induced cell death was observed in the two *sobir1*/*sobir1-like* knock-out mutant lines (middle two panels and lower two panels, respectively). C, Complementation by transient expression of *Nb*SOBIR1, *Sl*SOBIR1 or *Sl*SOBIR1-like, restores the Avr4/Cf-4-specific HR in the *N. benthamiana sobir1/sobir1-like* mutants, whereas this complementation does not take place upon transient expression of the corresponding kinase-dead mutants. *Nb*SOBIR1, *Sl*SOBIR1 or *Sl*SOBIR1-like, as well as their corresponding kinase-dead mutants (negative controls), were transiently co-expressed with Avr4/Cf-4, in the leaves of the *N. benthamiana sobir1/sobir1-like* knock-out lines. Each construct was agro-infiltrated at an optical density at 600 nm (OD_600_) of 0.5, and all leaves were photographed at 5 days post infiltration (dpi). Experiments were repeated at least three times, and similar results were obtained. Representative pictures are shown.

Over the years, in addition to Cf-4, more tomato RLPs have been identified that play a role in resistance to *C. fulvum*. An example is Cf-9, which confers recognition of the secreted *C. fulvum* effector Avr9 (van Kan et al., 1991; Jones et al., 1994). Transient co-expression of Cf proteins with their matching Avr ligands in *N. benthamiana* triggers a typical HR (Figure 1B) (van der Hoorn et al., 2000). Compared to the wild-type plant, none of the mutant lines was responsive to the Avr4/Cf-4 or Avr9/Cf-9 combination (Figure 1B), indicating that SOBIR1 and SOBIR1-like are indeed non-functional in the two mutant lines, due to disruption of their ORFs. Interestingly, complementation of the *SOBIR1(-like)* knockouts through transient expression of *Nb*SOBIR1, *Sl*SOBIR1, or *Sl*SOBIR1-like, together with the Avr4/Cf-4 combination, restored the HR (Figure 1C). Complementation did not take place upon co-expression of the corresponding kinase-dead mutants of SOBIR1, as in this case, the leaf tissue remained non-responsive to the Avr4/Cf-4 combination (Figure 1C). These results reinforce that SOBIR1/SOBIR1-like plays a pivotal role in RLP-mediated immunity and that the *sobir1*/*sobir1-like* mutant *N. benthamiana* plants form a robust basis for complementation studies.

We use *N. benthamiana*:*Cf-4* for the elucidation of the molecular mechanisms of signal transduction events triggered by Cf-4 upon Avr4 recognition (Liebrand et al., 2013; Wu et al., 2017; van der Burgh et al., 2019). Therefore, *SOBIR1* and *SOBIR1-like* were also knocked out in *N. benthamiana*:*Cf-4*. One homozygous mutant line was obtained with disruptions in both the ORF of *SOBIR1* and *SOBIR1-like*, which were introduced by Construct 1 (Figure 2A and Supplemental Figure S3). In agreement with our previous finding, transient expression of Avr4 triggered an HR in *N. benthamiana*:*Cf-4* plants, however, the knock-out line was non-responsive to Avr4 (Figure 2B), again confirming that Cf-4 functionality requires functional SOBIR1(-like). Complementation with *Nb*SOBIR1, *Sl*SOBIR1, or *Sl*SOBIR1-like, in combination with transient expression of Avr4, again resulted in a Cf-4-mediated HR, whereas complementation did not take place upon co-expression of the corresponding kinase-dead mutants of SOBIR1 (Figure 2C). Hereafter, this mutant line was further validated by monitoring the production of reactive oxygen species (ROS), which is a very early downstream response upon immune activation, upon treatment with either Avr4 or flg22. The latter is a peptide derived from bacterial flagellin that is perceived by the RLK FLAGELLIN-SENSITIVE 2 (FLS2) (Gòmez-Gòmez and Boller, 2000), which is also present in *N. benthamiana* and which does not interact with SOBIR1 and does not require SOBIR1 for its functionality (Liebrand et al., 2013; Albert et al., 2015). Unlike the rapid and monophasic ROS burst induced by the flg22 peptide, a biphasic ROS accumulation was observed when leaf discs of *N. benthamiana*:*Cf-4* were treated with Avr4 protein. In the latter case, the first transitory response was followed by a second, sustained ROS burst, which was of higher amplitude when compared to the initial ROS burst (Figure 2D). As expected, the biphasic ROS burst was completely abolished in the *N. benthamiana*:*Cf-4 sobir1/sobir1-like* mutant (Figure 2D), which further verifies that the mutant line has become non-responsive to Avr4. Intriguingly, an unexpected second sustained ROS burst, triggered by flg22, was observed in this *N. benthamiana*:*Cf-4 sobir1/sobir1-like* mutant, revealing that there is potential crosstalk taking place between RLP/SOBIR1- and FLS2-triggered pathways. We propose that the appearance of a biphasic ROS burst triggered by flg22 in the SOBIR1(-like) knock-out mutant, uncovers the presence of inhibitory activity of the RLP/SOBIR1 signal transduction pathway on the signaling route employed by FLS2.

**Figure 2.**
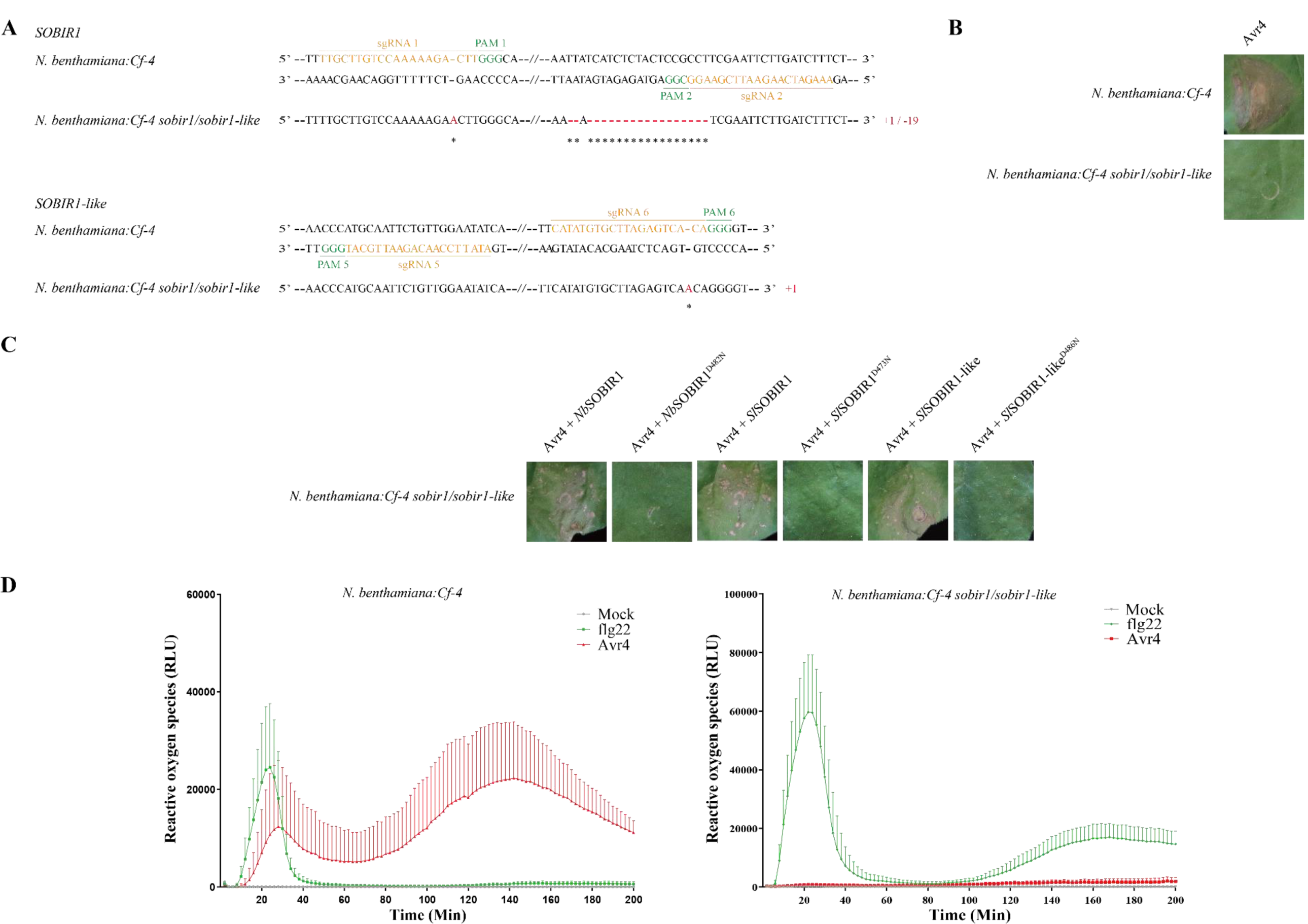
CRISPR/Cas9-induced targeted knockout of *SOBIR1* and *SOBIR1-like* in transgenic *N. benthamiana:Cf-4* abolishes the functionality of the *Cf-4* transgene. A, Nucleotide sequence alignment of the regions in *SOBIR1* (upper panel) and *SOBIR1-like* (lower panel) targeted by sgRNAs, with wild-type *SOBIR1* and *SOBIR1-like* sequences, respectively. The sgRNA sequences are shown in orange, and the PAM sites are indicated in green. The deleted nucleotides in the generated transformant are indicated with red dashes, and the inserted nucleotides are denoted with red letters. The type of mutations and the numbers of deleted/inserted nucleotides are shown on the right. B, *Agrobacterium*-mediated expression of Avr4 in *N. benthamiana:Cf-4* plants results in a rapid HR at the site of infiltration (upper panel), whereas agro-infiltration of Avr4 failed to induce cell death in the *N. benthamiana:Cf-4 sobir1*/*sobir1-like* knock-out line (lower panel). C, Complementation by transient expression of *Nb*SOBIR1, *Sl*SOBIR1 or *Sl*SOBIR1-like, restores the Avr4/Cf-4-specific HR in the *N. benthamiana:Cf-4 sobir1/sobir1-like* mutant, whereas this complementation does not take place upon transient expression of the corresponding kinase-dead mutants. *Nb*SOBIR1, *Sl*SOBIR1, or *Sl*SOBIR1-like, as well as their corresponding kinase-dead mutants (negative controls), were transiently co-expressed with Avr4, in the leaves of the *N. benthamiana:Cf-4 sobir1/sobir1-like* knock-out line. Each construct was agro-infiltrated at an OD_600_ of 0.5, and the leaves were photographed at 5 dpi. D, Avr4 fails to induce a reactive oxygen species (ROS) burst in the *N. benthamiana:Cf-4 sobir1*/*sobir1-like* knock-out line. Leaf discs of *N. benthamiana:Cf-4* (upper panel) and *N. benthamiana:Cf-4 sobir1*/*sobir1-like* (lower panel) were treated with 0.1 μM Avr4 or 0.1 μM flg22 (positive control), or with water (mock) (negative control). ROS production is expressed as relative light units (RLU), and the data are represented as mean + SD. Experiments were repeated at least three times, and similar results were obtained. Representative pictures are shown. Note that in the *N. benthamiana:Cf-4 sobir1*/*sobir1-like* knock-out line, the response to flg22 manifests itself as a biphasic ROS burst (lower panel), whereas in the non-mutant *N. benthamiana:Cf-4*, the flg22-triggered ROS burst is monophasic (upper panel).

## Supplemental Data

The following supplemental materials are available.

## Supplemental Methods

**Supplemental Figure S1**. Nucleotide sequence of NbSOBIR1 and NbSOBIR1-like.

**Supplemental Figure S2**. Determination of the effectiveness and efficiency of the generated CRISPR/Cas9 constructs.

**Supplemental Figure S3**. Phenotypes of wild-type *N. benthamiana* and the various mutant lines.

**Supplemental Table S1**. Sequences of the six sgRNAs.

**Supplemental Table S2**. Sequences of the primers used in this study.

## ACKNOWLEDGMENTS

We thank Bert Essenstam and Henk Smid from Unifarm for excellent plant care.

