## Supplemental Materials for "Knocking out *SOBIR1* in *Nicotiana benthamiana* abolishes functionality of transgenic receptor-like protein Cf-4"

### Supplemental Methods

#### CRISPR/Cas9 constructs generation and testing

To knock out functional *SOBIR1* and *SOBIR1-like* in *N. benthamiana*, two constructs were generated using the Golden Gate cloning method (Engler et al., 2014). Both constructs contain four sgRNAs. In total, six different sgRNAs were designed, with sgRNA1/2/3/4 targeting different regions of the open reading frame (ORF) of *SOBIR1*, and sgRNA5/6 targeting the ORF of *SOBIR1-like* (Supplemental Table S1 and Figure S1). Each oligonucleotide pair for generating the different sgRNAs was annealed into double-stranded DNA (Supplemental Table S2), followed by assembly into the Level 0 vector pICSL01009:AtU6p (SOL7857). Hereafter, each sgRNA cassette was inserted into the appropriate Level 1 vectors (SOL7861, SOL7862, SOL7863 and SOL7864), whereby pICH47751::AtU6p::sgRNA1 (SOL7867), pICH47761::AtU6p::sgRNA2 (SOL7868), pICH47751::AtU6p::sgRNA3 (SOL7869), pICH47761::AtU6p::sgRNA4 (SOL7870), pICH47772::AtU6p::sgRNA5 (SOL7871) and pICH47781::AtU6p::sgRNA6 (SOL7872), were obtained. Together with pICH47732::NOSp-BAR-NOST (SOL7859), pICH47742::2×35S::hCas9 (SOL7860), pICH41822 end-linker 6 (SOL7865), pICH47772::AtU6p::sgRNA5 and pICH47781::AtU6p::sgRNA6, pICH47751::AtU6p::sgRNA1 and pICH47761::AtU6p::sgRNA2 were assembled into the Level 2 vector pAGM4723 (SOL7866), resulting in CRISPR/Cas9 Construct 1; pAGM4723::BAR::Cas9::sgRNA1:: sgRNA2:: sgRNA5:: sgRNA6 (SOL7880). Next to this, pICH47751::AtU6p::sgRNA3 and pICH47761::AtU6p::sgRNA4 were also assembled into the Level 2 vector pAGM4723, resulting in CRISPR/Cas9 Construct 2; pAGM4723::BAR::Cas9::sgRNA3:: sgRNA4:: sgRNA5:: sgRNA6 (SOL7881).

Each construct was transformed individually into *Agrobacterium tumefaciens* strain GV3101, followed by agro-infiltration at an optical density at 600 nm ($OD_{600}$) of 1 in leaves of *N. benthamiana* as described before (van der Hoorn et al., 2000). Genomic DNA of the agro-infiltrated leaves was isolated at 5 days post infiltration (dpi) (Fulton et al., 1995), and the targeted fragments of *SOBIR1* and *SOBIR1-like* were amplified, and of which the *SOBIR1-like* fragments were digested with *HpyCH*4V and *Hinf*I at 37 °C for 1 h (New England Biolabs) (Supplemental Table S2 and Figure S2).

#### Plant transformations and mutation detection

*Agrobacterium* carrying the different CRISPR/Cas9 constructs was used for transformation into wild-type *N. benthamiana* and *N. benthamiana:Cf-4* (Gabriëls et al., 2006), respectively (Rorsch et al., 1988). To select homozygous transformants and to determine the mutation type, genomic DNA was extracted from each transformant (Fulton et al., 1995) and used as a template to amplify the targeted fragments of *SOBIR1* and *SOBIR1-like* with the specific primers listed in Supplemental Table S2. Hereafter, the purified fragments were analyzed by Sanger sequencing.

#### Plant growth conditions

*N. benthamiana*, *N. benthamiana:Cf-4* and all generated transformants of these plants were grown in a climate chamber under 15 h of light at 21 °C and 9 h of darkness at 19 °C, with a relative humidity of ~70%.

#### *Agrobacterium*-mediated transient transformation

*Agrobacterium*-mediated transient transformation of Avr4, the Avr4/Cf-4 combination, or the Avr9/Cf-9 combination, in five-week-old *N. benthamiana* plants, was performed as described before ($OD_{600}$ = 0.5 for each construct) (van der Hoorn et al., 2000). For complementation studies, eGFP-tagged *Nb*SOBIR1 (SOL2911), *Sl*SOBIR1 (SOL2774) or *Sl*SOBIR1-like (SOL2773), as well as their corresponding kinase-dead mutants *Nb*SOBIR1$^{D482N}$ (SOL7928), *Sl*SOBIR1$^{D473N}$ (SOL2875) or *Sl*SOBIR1-like$^{D486N}$ (SOL2876) (Liebrand et al., 2013), were mixed with Avr4 ($OD_{600}$ = 0.8 for each construct), and then syringe-infiltrated into fully expanded leaves of *N. benthamiana:Cf-4 sobir1/sobir1-like*. The development of an HR was photographed at 5 dpi.

#### Reactive oxygen species (ROS) assay

ROS production was determined by a luminol-based assay (Keppler et al., 1989). For this, leaf discs from 4- to 5-week-old *N. benthamiana:Cf-4* and *N. benthamiana:Cf-4 sobir1/sobir1-like* plants were collected by using biopsy punches (Ø 5 mm, Robbins Instruments) and incubated in 100 µL of sterile water in a 96-wells plate overnight. The water was then replaced carefully by 50 µL of fresh sterile water and the leaf discs were incubated for another hour. 50 µL of a freshly prepared reaction solution, containing 100 µM of luminol (L-012, Fujifilm, Japan), 20 µg/mL horseradish peroxidase (Sigma), and 0.2 µM flg22 (EZBiolab) or 0.2 µM Avr4, was added to each well. For the mock treatment, no flg22 or Avr4 was added. Chemiluminescence was monitored using a CLARIOstar plate reader (BMG Labtech), with a program of 100 cycles and 2 min per cycle.

**A**

ATGGCCTTCACTGCTTCACAAATTCACTTCTTTTTCTTCTCACTTTTCGCCTTTTTACTTATTGCTGTTCAAGCAAGACTGAAT
CTTTATCCACCAGATCATGCTGCACTTTTGCTTGTCCAAAAAGACTTGGGCATCCAAGGTCAACGCATTGCACTTTGCAACT
CTGCAACAATATCCTGTGAAAGGCGAAAGGCAAACAGAACACAATTGTTGAGAGTCACCCGTATTGACTTCAGATCCAGTG
GATTGAGTGGAACTTTATCTCCTGCCATTGGAAAACTTTCTGTGCTCAAAGAACTCTCTCTTCCAAACAACCAACTCTTTGAC
CAAATCCCAGTTCAGATTCTTGATTGCCGTAAACTGGAGATTCTTGACCTTGGAAACAATCTGTTTTCTGGGAAAGTTCCATC
TGAATTATCATCTCTACTCCGCCTTCGAATTCTTGATCTTTCTTCAAATGAGTTTTCTGGGAATCTTAACTTCTTGAAGTATTT
TCCCAACTTAGAAAAACTCTCTTTAGCTGATAACATGTTCACTGGAAAAATACCCCCTTCTTTGAAATCTTTTAGGAATCTCC
GTATCCTTAATATTTCAGGAAACAGTTTTCTTGAAGGTCATGTGCCTGTTATGAGTCAAGTTGAGCACTTGTCAGCAGAATT
GGATCAGCACTTCGTTCCAAAACGTTACATTCTTGCTGAAAATTCAACAAGGTCAAATCAGATATCAGCACTGGCACCTAAT
TCCAATTCAGGAAATGCCCCAGCTCCAGCACCGAGTCATAATGTCACTCCAATCCATAAACATAGTAACAGGAAGAAAAGG
AAAGTCAGAGCGTGGTTACTTGGTTTCTTTGCTGGTTCTTTCGCAGGAGCTATATCTGCAGTACTCTTATCAGTTCTTTTTAA
GCTGGTCATGTTTTTTGTCCGAAAGGGAAAGACTGATGGAACTTTAACAATATACAGTCCACTGATTAAGAAAGCCGAGGAT
TTGGCCTTCTTAGAGACAGAAGATGGAGTAGCATCACTTGAAATGATTGGAAAAGGTGGATGCGGAGAAGTTTATAGAGCT
GAGTTACCGGGAAGTAATGGGAAGATTATAGCTATAAAGAAGATTATACAACCCCCAATGGATGCTGCAGAACTCACCGAG
GAAGATACCAAGGCTTTGAACAAAAAAATGCGACAAGTAAAATCAGAAATTCAAATTCTTGGTCAAATCAGACACAGGAATC
TGCTTCCCCTACTGGCACATATGCCTAGGCCAGACTGCCATTACTTGGTATATGAATATATGAAAAATGGGAGCTTACAAGA
TATCCTCCAGCAAGTCACAGAAGGGACAAGGGAATTAGATTGGTTGGGACGACACCGAATTGCAGTAGGGATAGCTTCTG
GACTTGAGTATCTCCATATAAACCACAGTCAATGCATAATTCACAGAGATCTAAAGCCAGCAAATGTCCTTCTTGACGATGA
TATGGAAGCTCGAATTGCTGATTTTGGACTTGCAAAAGCACTCCCAGATGCCCATACACATGTTACGACTTCAAATGTTGCA
GGAACTGTGGGATATATTGCACCAGAATACCATCAGACACTGAAGTTTACGGGTAAGTGTGATATATACAGCTTTGGTGTG
GTGTTGGCAGTGCTGGTTATAGGAAAACTTCCATCAGATGAATTTTTCCAGCACACGCCTGAGATGAGTTTAGTGAAGTGG
CTGAGAAATGTAATGACTTCTGAGGATCCGAAAAGGGCAATTGATTCAAAGCTGATAGGAAATGGATTTGAGGAGCAAATG
CTTTTGGTTCTCAAGATAGCTTGCTTTTGTACTCTGGAGAATCCAAAGGAGAGGCCTAACAGTAAGGATGTTAGGTGTATGT
TAACTCAGATCAAGCATTAG

**B**

ATGGCCTCCACTACCTCCCATATTCACCTATCTCTTCTGTCTCTCTTCACTCTTATCCTTCATGTTCAAGCAAGAATCAACCTT
TACTCGCCAGATTATAGTGCTCTTTTGGTTGTCCGAAAAGGCTTAGACGTCCCTAGACAACTTAGTGCTATAGAGAACCCAT
GCAATTCTGTTGGAATATCATGTGAACGACGACTCACAAACAATTCATATGTGCTTAGAGTCACAGGGGTTGTTTTCAAGTC
CTATGAACTGAGGGCAACTCTATCTCCTGCCATTTGCAAGCTTTCTGAGTTCAAAAAATTGTCCCTACAAAACAACAAGCTCT
TTGACAGAATCCCAACTCAAATTGTTGAGGTGCCGAAAATTGGAAATCTTGAACCTTCAAAACAACCAGTTTTCTGGCAAAG
TCCCATCTGAATTATCATCTCTTGTCCGCCTTCGAATCCTTGACCTCTCTTCTAATGAATTATCAGGGAACCTCAATTTCTTGA
AATACTTTCCCAACCTTGAAAATTTGTCCCTTGCTGATAACATGTTCATTGACAAAATACCTCAATCCTTGAAATCTTTCAGAA
ATCTCCGGTTCCTCAACATTTCAGGGAATAGTTTTCTTTAAGGTCCGGTGCCTTTCATGAGTCAAGTTAAGCACTTATCAGCA
GACTTGAATCGAAAAGATTATGTTCTTAAACGTTACATTCTTGCTGAAAACTCAACAAGGTCAAGTCACAAATCTGCAATGG
CACCACAGTCCTATTCAGGAAATGCCCCGGCTCCAGGACCCAGTGCAATTGTAGTACCAGTACACAAACATAACAACACAA
AAAAGAAGGTAAGCATATGGATTCTTGGATTCCTGGCTGGATCTTTTGCAGGGATCTTATCTGCGTGGTCTTATCCTTGCTC
TTCAAGGTGGTTATGATTATCGGAGGGAGCAAGGATGATTCAGGTATAACAATTTTCAGTCCATTGATTAAGAAAGCAGAGT
TACCAGGAAGTAATGGAAAGATTATTGCTATAAAGAAGATCGTAGAACCCCCAAAGGATGCTGCAGAACTCACTGAGGAAG
ATAGCAAGGCTTTGAATATTAAAATGTGCCAGACCAAATCAGAAATTAAAATTGTAGGTCAAATAAGACACAGGAATTTGCT
TCCTCTAATAGCACATATGCCAAGACCTGACTGCCATTACTTGGTCCGTGAGTACATGAAAAATGGGAGCTTACAGGATGCC
GTCCAGCAAGTCACGGAGGGGACACGGGAAATAGATTGGTCGGCACGTCATCGAATTGCAGTCGGGATCGCTGCTGGGCT
TGAGTATCTTCATGTAAATCATAGTCAGCGTACAATTCACAGGGATCTAAAGCCAGCAAATGTCCTTCTTGCGAAAGCAATC
CCAGAATCCCTTACGCTTTTCAACTTCACACGTGGTAGGAACTTTAGGATACATTGCACCAGAATATTACCAGACCGTTAAG
TTCACAGATAAGTGTGATATATACAGCTTCGGGATGCTGCTAGGCGTGCTAGTTATGGGAAAGTTTCCCTCTGATGAGCTCT
TCCAGCCGGTTTCTGGGATGGGTTTAGTGAAATGGATGAGAAATGTCACGACTTCTGAGAATCCAAAAAGAGCAATTGATC
CAAAGCTGATGGGTAATGGGTATGAGGAGCAAAATGCTTTTGGTTCTCAAGATTGCCTGCTTTTGCTCACTGGATAATGCAA
AGGAGAGGCCTAACAGTAAGGATGTTAGATGCATGTTAACTCAGATCAAGCCTCAAGTAGGTAGAGGATAA

**Supplemental Figure S1.** Nucleotide sequence of *NbSOBIR1* and *NbSOBIR1-like*. The primers used for amplifying and sequencing targeted fragments of A, *SOBIR1* and B, *SOBIR1-like* are indicated in yellow. single-guide RNAs (sgRNAs) are highlighted in orange, followed by protospacer associated motifs (PAMs) that are shown in green.

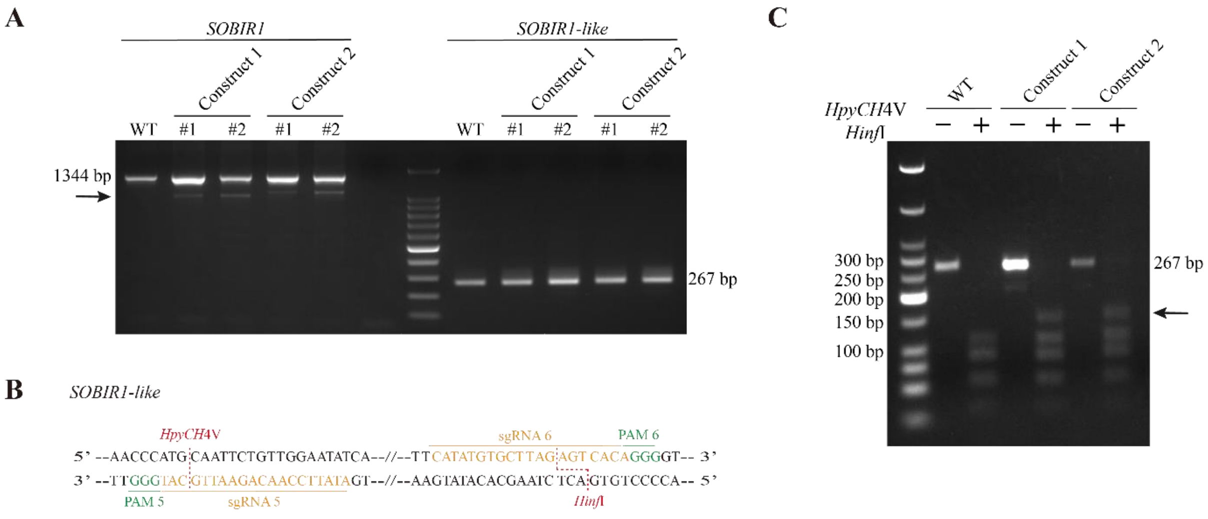


**Supplemental Figure S2.** Determination of the effectiveness and efficiency of the generated CRISPR/Cas9 constructs. A, PCR-based detection of CRISPR/Cas9-induced deletions in *NbSOBIR1* and *NbSOBIR1-like*. The fragments of *NbSOBIR1* (shown on the left) were amplified by PCR with the primer pair Nbsobir1fwd/Nbsobir1rev, while the fragments of *SOBIR1-like* (shown on the right) were amplified with the primer pair Nbsobir1likefwd/Nbsobir1likerev. Compared to wild type (WT), transient expression of Construct 1- and 2- induced the expected deletions (black arrow) in *SOBIR1*, indicating the high editing efficiency of Construct 1 in sgRNA1- and sgRNA2-targeted regions, and of Construct 2 in sgRNA3- and sgRNA4-targeted regions. However, no deletion was observed in *SOBIR1-like*. B, Nucleotide sequence alignment of the regions in *SOBIR1-like* targeted by sgRNA5 and sgRNA6. The sgRNA sequences are indicated in orange, the PAMs are shown in green, and the two restriction sites (*HpyCH*4V and *Hinf*I) in the sgRNAs are denoted in red. C, Digestion of the *SOBIR1-like* amplicons with *HpyCH*4V and *Hinf*I. *SOBIR1-like* amplicons were purified and digested with the two restriction enzymes, compared to WT an extra band (black arrow) was observed in Construct 1- and 2-treated *SOBIR1-like*, indicating that at least one restriction site was missing. Therefore, both constructs were also able to edit the sequence of *SOBIR1-like*.

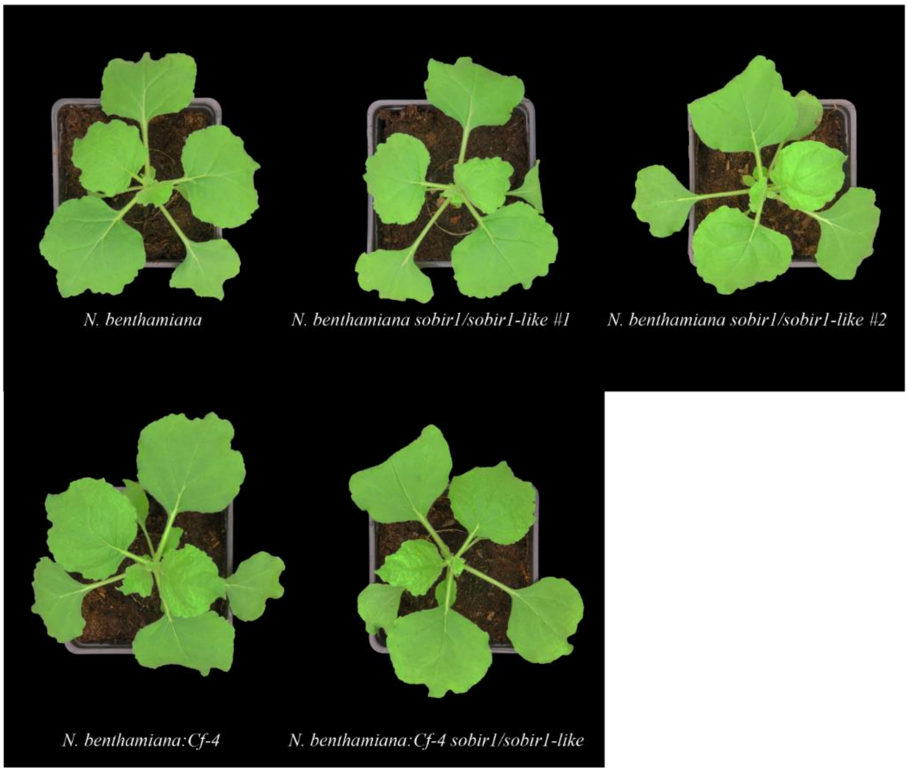


**Supplemental Figure S3.** Phenotypes of wild-type *N. benthamiana* and the various mutant lines. All plants were grown on soil and photographed when four to five-weeks-old. Note that the mutant lines do not have an obvious phenotype.

**Supplemental Table S1.** Sequences of the six sgRNAs.

| | Sequence (5′-3′) | Target gene |
|---|---|---|
| sgRNA1 | TTGCTTGTCCAAAAAGACTT | *SOBIR1* |
| sgRNA2 | AAAGATCAAGAATTCGAAGG | *SOBIR1* |
| sgRNA3 | TCCACTGATTAAGAAAGCCG | *SOBIR1* |
| sgRNA4 | CTAGGCATATGTGCCAGTAG | *SOBIR1* |
| sgRNA5 | ATATTCCAACAGAATTGCAT | *SOBIR1-like* |
| sgRNA6 | CATATGTGCTTAGAGTCACA | *SOBIR1-like* |

**Supplemental Table S2.** Sequences of the primers used in this study.

| Primer code | Primer name | Sequence (5′-3′) | Note |
|---|---|---|---|
| ho51 | sgRNA1_fw | ATTGTTGCTTGTCCAAAAAGACTT | sgRNA1 |
| ho52 | sgRNA1_rev | AAACAAGTCTTTTTGGACAAGCAA | |
| ho53 | sgRNA2_fw | ATTGAAAGATCAAGAATTCGAAGG | sgRNA2 |
| ho54 | sgRNA2_rev | AAACCCTTCGAATTCTTGATCTTT | |
| ho55 | sgRNA3_fw | ATTGTCCACTGATTAAGAAAGCCG | sgRNA3 |
| ho56 | sgRNA3_rev | AAACCGGCTTTCTTAATCAGTGGA | |
| ho57 | sgRNA4_fw | ATTGCTAGGCATATGTGCCAGTAG | sgRNA4 |
| ho58 | sgRNA4_rev | AAACCTACTGGCACATATGCCTAG | |
| ho59 | sgRNA5_fw | ATTGATATTCCAACAGAGTTGCAT | sgRNA5 |
| ho60 | sgRNA5_rev | AAACATGCAACTCTGTTGGAATAT | |
| ho61 | sgRNA6_fw | ATTGCATATGTGCTTAGAGTCACA | sgRNA6 |
| ho62 | sgRNA6_rev | AAACTGTGACTCTAAGCACATATG | |
| ho11 | Nbsobir1fwd | ATGGCCTTCACTGCTTCACAAAT | To amplify *SOBIR1* and test the effectiveness of the constructs. |
| ho12 | Nbsobir1rev | CTGGAGGATATCTTGTAAGCTCCC | |
| ho13 | Nbsobir1likefwd | CCTTTACTCGCCAGATTATAGT | To amplify *SOBIR1-like* and test the effectiveness of the constructs. |
| ho14 | Nbsobir1likerev | TGAGTTGGGATTCTGTCAAAG | |
| ho78 | NbSOBIR1 1-2 seq1_fw | ACTTTTCGCCTTTTTACTTATTGC | To amplify and sequence *SOBIR1*-containing sgRNA1&2 region. |
| ho79 | NbSOBIR1 1-2 seq2_rev | TGGAACGAAGTGCTGATCCAAT | |
| ho80 | NbSOBIR1 2-2 seq2_fw | ATTGGATCAGCACTTCGTTCC | To amplify and sequence *SOBIR1*-containing sgRNA3&4 region. |
| ho81 | NbSOBIR1 2-2 seq1_rev | AATTCGAGCTTCCATATCATCGT | |
| ho82 | NbSOBIR1-like seq1_fw | CTATCTCTTCTGTCTCTCTTCAC | To amplify and sequence *SOBIR1-like*-containing sgRNA5&6 region. |
| ho83 | NbSOBIR1-like seq4_rev | CATAATCTTTTCGATTCAAG | |
